# Large increases in resistance training volume do not impair skeletal muscle hypertrophy or anabolic–catabolic molecular signalling in trained individuals

**DOI:** 10.64898/2026.02.23.707462

**Authors:** Júlio B. B. Camargo, Diego Bittencourt, J. Max Michel, Deivid G. Silva, João G. A. Bergamasco, Dakota R. Tiede, Dustyn Lewis, Enya T. A. Nacafucasaco, Otávio Ferrari, Ana C. C. Melo, Matheus Iasulaitis, Marcelo Rebelato, Michael D. Roberts, Cleiton A. Libardi

**Affiliations:** MUSCULAB – Laboratory of Neuromuscular Adaptations to Resistance Training, Department of Physical Education, Federal University of São Carlos (UFSCar), São Carlos, SP, Brazil; School of Kinesiology, Auburn University, Auburn, AL, USA; Healthspan, Resilience and Performance Research, Florida Institute for Human and Machine Cognition, Pensacola, FL, USA

**Keywords:** muscle size, training variables, weekly sets, protein turnover

## Abstract

Skeletal muscle hypertrophy results from the integrated regulation of anabolic and proteolytic processes in response to mechanical loading. Although increases in resistance training (RT) volume are used to increase mechanical stress, it remains uncertain whether large and abrupt volume progressions could exceed muscle adaptive capacity by disrupting the balance between anabolic and catabolic signaling. The present study investigated whether a large increase in weekly RT volume (+120%) leads to impaired hypertrophic outcomes and intracellular regulatory responses compared with a modest increase (+20%). Twenty-five resistance-trained men and women (18–35 years old) completed an 8-week randomized, single-blind, within-subject unilateral intervention. Each participant trained both legs twice weekly, with one leg assigned to the large (VOL120) and the contralateral leg to the modest (VOL20) weekly volume progressions relative to habitual training volume. Vastus lateralis muscle cross-sectional area (mCSA) was assessed by ultrasonography before and after training. Muscle biopsies were obtained at baseline, post-intervention, and 24 h after the last session to quantify muscle fiber cross-sectional area (fCSA), satellite cell myonuclear content, and anabolic/catabolic signaling markers. Both protocols induced increases in mCSA over time (*p*<0.001), with no protocol vs. time interaction. No significant effects were observed for fCSA nor satellite cell number or myonuclear content. Additionally, molecular responses related to translational regulation and protein degradation were largely similar between protocols. Collectively, these data indicate that a large, abrupt increase in weekly set volume does not impair hypertrophic adaptations or meaningfully alter the anabolic–catabolic signaling profile in resistance-trained individuals.

## INTRODUCTION

Skeletal muscle hypertrophy represents a complex adaptive process that emerges from the coordinated regulation of protein synthesis, protein degradation, and structural remodeling in response to repeated mechanical loading (Roberts, McCarthy et al. 2023). While the intrinsic regulatory capacity underlying such processes ultimately constrains the magnitude and sustainability of hypertrophic adaptations (Roberts, McCarthy et al. 2023), progressive increases in loading during resistance training (RT) are commonly implemented to maintain adaptive responses across prolonged training periods (ACSM 2009, Schoenfeld, Fisher et al. 2021). Among the variables used to manipulate loading, RT volume represents a practical means of experimentally increasing cumulative mechanical stress, particularly in resistance-trained individuals (Israetel, Feather et al. 2020). Consistent with this rationale, experimental and meta-analytical evidence has frequently reported greater hypertrophy following higher-volume RT protocols, supporting a dose-dependent relationship between volume and muscle growth (Burd, Holwerda et al. 2010, Schoenfeld, Ogborn et al. 2017, Schoenfeld, Contreras et al. 2019, Brigatto, Lima et al. 2022). More recently, meta-regression analyses suggest diminishing returns with increasing weekly set volume, yet whether an upper boundary exists beyond which additional volume attenuates further hypertrophic adaptation remains unresolved, particularly in resistance-trained individuals (Pelland, Remmert et al. 2025). Notably, when examining studies conducted exclusively in resistance-trained participants, several investigations have reported comparable hypertrophic adaptations despite substantial differences in prescribed RT volume (Heaselgrave, Blacker et al. 2019, Aube, Wadhi et al. 2022, Enes, EO et al. 2024, Moreno, Sampson et al. 2024). These discrepant findings raise the possibility that abrupt, high-magnitude increases in training volume (i.e., implemented all at once rather than progressively) may challenge the muscle’s adaptive capacity and attenuate hypertrophic responses rather than enhance them (Camargo, Bittencourt et al. 2025).

Understanding these divergent morphological outcomes requires integrating whole-muscle responses with the intracellular pathways that regulate translational efficiency, translational capacity, and protein breakdown. Activation of the mammalian target of rapamycin complex 1 (mTORC1) regulates translational efficiency and myofibrillar protein synthesis through downstream targets such as p70S6K and 4E-BP1 (Roberts, McCarthy et al. 2023). In parallel, ribosome biogenesis also contributes to translational capacity (Bamman, Roberts et al. 2018), while satellite cell–mediated myonuclear accretion supports sustained fiber growth by expanding the cellular machinery required for elevated protein synthesis (Petrella, Kim et al. 2008, Conceicao, Vechin et al. 2018). Indeed, training volume has been shown to modulate several of these processes, including satellite cell responses and ribosome biogenesis (Hanssen, Kvamme et al. 2013). In contrast, the effects of chronic, progressive increases in RT volume on canonical anabolic signaling pathways remain insufficiently characterized in resistance-trained humans, as most available evidence derives from acute studies employing relatively modest exercise volumes (Burd, Holwerda et al. 2010, Terzis, Spengos et al. 2010, Kumar, Atherton et al. 2012, Ahtiainen, Walker et al. 2015). Hypertrophic adaptations are also influenced by the regulation of muscle protein breakdown, which contributes to net protein balance and structural remodeling. Calpains and the ubiquitin–proteasome system play complementary roles in the degradation of damaged or misfolded proteins and cytoskeletal turnover in skeletal muscle (Sandri 2013, Hyatt and Powers 2020). Experimental evidence from animal models indicates that excessive mechanical loading or insufficient recovery between resistance exercise bouts can sustain proteolytic signaling without proportional increases in protein synthesis, thereby impairing hypertrophic adaptations (Takegaki, Ogasawara et al. 2017, Takegaki, Ogasawara et al. 2019). In this sense, it is possible that large and abrupt increases in RT volume shift the balance between anabolic and catabolic processes toward a proteolytic-dominant state, effectively uncoupling intracellular signaling responses from morphological adaptation.

Collectively, these observations suggest a non-linear relationship between RT volume and muscle hypertrophy, whereby volume-dependent gains occur up to a threshold beyond which further increases in mechanical stress exceed the adaptive capacity of skeletal muscle. Under this framework, moderate increases in volume may optimize adaptive signaling, whereas excessive volumes may compromise net protein balance and tissue remodeling, consistent with an inverted “U”-shaped volume–hypertrophy relationship (Figueiredo, de Salles et al. 2018, Camargo, Bittencourt et al. 2025). Evidence consistent with this model has emerged from individual-level analyses in resistance-trained humans. Scarpelli et al. (2022) reported smaller increases in vastus lateralis muscle cross-sectional area (mCSA) in limbs exposed to large volume progressions (83–120% above baseline) compared with contralateral limbs subjected to a modest increase (∼20%). Accordingly, Haun et al. (2018) reported that, in resistance-trained men (n = 31), large weekly increases in back-squat volume over 6 weeks resulted in no increases in vastus lateralis muscle thickness or type I and type II fiber cross-sectional area (fCSA). However, findings from both studies were not derived from a randomized controlled design specifically intended to test whether large, abrupt increases in RT volume directly attenuate hypertrophy or to determine whether such effects are accompanied by coordinated alterations in anabolic and proteolytic signaling pathways.

Therefore, the primary aim of the present study was to determine whether a large and abrupt increase in weekly RT volume (+120%; VOL120) attenuates hypertrophic adaptations at both the whole-muscle and muscle fiber levels compared with a protocol involving a modest volume increase (+20%; VOL20) in resistance-trained individuals. Additionally, we examined acute and chronic responses of key anabolic and proteolytic biomarkers under these distinct volume-progression protocols. We hypothesized that the larger volume progression would exceed muscle adaptive capacity, leading to attenuated hypertrophy and molecular responses consistent with a shift toward catabolic regulation relative to the modest progression.

## METHODS

### Ethical approval and participants

Healthy young men and women aged 18–35 years were recruited. Eligibility criteria included a minimum of 1 year and a maximum of 5 years of consistent lower-limb RT experience, with a habitual quadriceps training volume ranging from 12 to 20 sets per week, including leg press and leg extension exercises. Participants were excluded if they reported chronic use of anabolic steroids, vitamin or ergogenic supplements, or anti-inflammatory medications, or if they had any musculoskeletal injury or neuromuscular disorder that could interfere with participation in the prescribed training protocols. All participants provided written informed consent after receiving a detailed explanation of the study procedures, risks, and benefits. The study was approved by the Institutional Review Board (approval no. 6.642.786), conducted in accordance with the Declaration of Helsinki, and registered in the Brazilian Registry of Clinical Trials (RBR-2f836zh).

### Experimental design

This study employed a randomized, single-blind, within-subject unilateral design with repeated measures. This approach was selected to minimize inter-individual variability, allow each participant to serve as their own control, and increase statistical sensitivity for detecting training-induced adaptations while reducing the required sample size (MacInnis et al., 2017). Each lower limb represented an independent experimental unit and was randomly allocated to one of two resistance training (RT) volume-progression protocols: a modest increase in weekly volume (+20%; VOL20) or a large, abrupt increase (+120%; VOL120) both relative to each participant’s habitual pre-study weekly set volume, and maintained throughout the 8-week training period (16 sessions). The primary outcome was vastus lateralis mCSA. Secondary outcomes included muscle fibre morphology, satellite cell and myonuclear content, and molecular markers associated with anabolic and proteolytic pathways.

Figure 1 provides an overview of the experimental timeline. Briefly, vastus lateralis mCSA was assessed by ultrasonography at baseline and then reassessed 72 h later (pre-biopsy) to determine the typical error and coefficient of variation of the ultrasound method. After completion of the training intervention, mCSA was reassessed 96 h after the final training session, followed by bilateral muscle biopsies to evaluate chronic adaptations (Pre vs. Post). Participants then completed an additional training session (session 17), after which muscle biopsies were obtained 24 h later to assess acute molecular responses (0 h vs. 24 h). During this final session, each limb performed the same RT protocol to which it had been allocated throughout the intervention.

**Figure 1.**
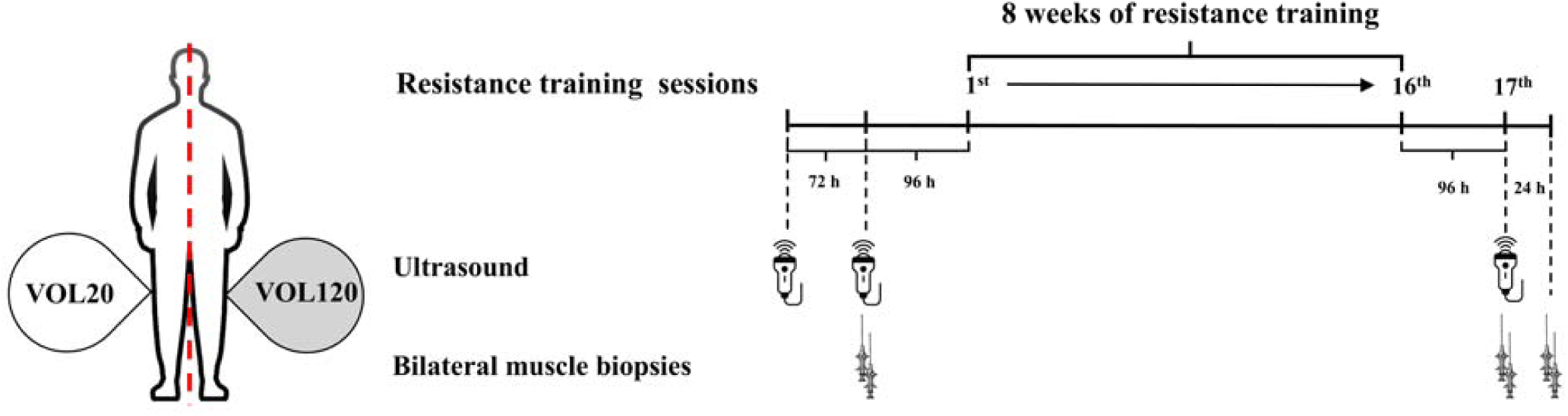
Experimental design. VOL20: modest increase in weekly resistance training volume (+20%); VOL120: large increase in weekly resistance training volume (+120%).

### Sample size estimation

Sample size estimation was performed using a simulation-based power analysis (10,000 simulations) for a two-way repeated-measures design, implemented with the Superpower package in R (Lakens and Caldwell 2021). Statistical power was set at 80% with a two-sided α level of 0.05. Based on previous unilateral training studies, a correlation coefficient of *r* = 0.78 was assumed for changes in vastus lateralis muscle mCSA between limbs. Although prior work investigating training volume reported effect sizes of approximately *d* = 0.75, a more conservative effect size (*d* = 0.50) was adopted to account for differences in experimental design. Accordingly, a Cohen’s *f* value of 0.20 was assumed for the interaction effect. Under these assumptions, the estimated minimum sample size was 24 participants. To account for potential attrition, recruitment exceeded the minimum number required by the power analysis.

### Resistance training protocols

Participants completed two supervised RT sessions per week for 8 weeks, with all sets performed to concentric muscle failure within a target repetition range of 9–12 repetitions and a fixed 2-min inter-set rest interval. Training loads were adjusted on a set-by-set basis: loads were increased if more than 12 repetitions were completed and reduced if fewer than 9 repetitions were achieved. Both RT protocols were individualized based on each participant’s habitual quadriceps training volume prior to study entry. Baseline weekly volume was determined via structured interviews in which participants reported the number of exercises, sets per exercise, and weekly training frequency for the quadriceps muscles. Weekly volume was calculated as the product of exercises per session, sets per exercise, and weekly frequency. Based on this value, each limb was assigned to either a modest volume increase (+20%; VOL20) or a large, abrupt volume increase (+120%; VOL120), which was maintained throughout the intervention period. The exercises performed in each session were the 45° leg press and the leg extension (Effort NKR; Nakagym, São Paulo, Brazil). Both protocols were performed concurrently within the same participant, with each leg assigned to one training protocol. Limb allocation was randomized using block randomization stratified by limb dominance (right/left). The exercise order was kept constant across all sessions (45° leg press followed by leg extension). To minimize potential order effects between protocols, the limb corresponding to each protocol alternated as the first limb trained across sessions. Volume load (load × sets × repetitions) was recorded for all sessions.

Participants were instructed to refrain from any unsupervised quadriceps resistance training or high-intensity lower-limb exercise throughout the study period. Unsupervised training targeting the upper body, gluteal, hamstring, and calf muscles was permitted, with participants explicitly instructed to avoid exercises likely to substantially load the quadriceps (e.g., squat variations, lunges, deadlifts).

### Dietary adherence

To minimize dietary confounding, participants were instructed to maintain their habitual dietary intake throughout the study and to refrain from using nutritional supplements other than that provided by the researchers. Immediately after each RT session throughout the intervention, participants ingested 30 g of whey protein (Adaptogen®, São Paulo, Brazil) to standardize post-exercise protein intake and reduce between-participant variability in the acute muscle protein synthetic response to exercise (Tipton, Ferrando et al. 1999, Hulmi, Lockwood et al. 2010, Witard, Jackman et al. 2014). Dietary adherence was assessed by a self-reported 24-h dietary records using MyFitnessPal software (http://www.myfitnesspal.com), which presents good relative validity for tracking energy and macronutrient consumption (Teixeira, Voci et al. 2018). Dietary recall was collected before each of the 3 muscle biopsy samples. Participants were instructed on how to properly report all food items and their respective portion sizes consumed for each meal. An experienced researcher inserted each food item into the software, which provided information regarding total energy consumption, and total grams for all macronutrients (i.e., proteins, fats, and carbohydrates).

### Ultrasound assessment

Vastus lateralis mCSA was obtained through B-mode ultrasound (US) images using a 7.5 MHz linear probe (MySono U6, Samsung), following the procedures validated by Lixandrão et al. (Lixandrao, Ugrinowitsch et al. 2014). Participants were instructed to refrain from vigorous physical activity for at least 72 h before each assessment. Before the assessments, participants laid down for 15 minutes to ensure fluid redistribution. Then, the assessor measured the distance between the greater trochanter and the lateral epicondyle of the femur to determine the length of the right and left femur. The point corresponding to 50% of the length was marked with a dermographic pen as a reference point for image acquisition. From the 50% point, the skin was marked in the medial and lateral directions every 2 cm to guide the displacement of the probe in the sagittal plane, parallel to the long axis of the femur. Water-soluble transmission gel was applied to ensure acoustic coupling of the US probe without compressing the epidermis. To obtain the mCSA, the US probe was displaced in the sagittal plane, starting at the alignment point of the upper edge of the probe with the most medial skin mark (over the rectus femoris muscle) and ending on the lateral side of the thigh. Then, the images were opened in the software Power Point (Microsoft, USA) and manually rotated to reconstruct the entire fascia of the vastus lateralis muscle to allow panoramic visualization of the entire mCSA (Reeves, Maganaris et al. 2004). The reconstructed image was opened in ImageJ software, and the “polygonal” function was used to circle and measure the CSA of the vastus lateralis, excluding the underlying muscle fascia and bone tissue as much as possible. The typical error and the coefficient of variation were 0.68 cm² and 2.31%, respectively. Fascicle length and pennation angle of vastus lateralis were measured at the same time and site of the mCSA acquisition, with the probe placed longitudinally to the muscle belly. The PA was defined as the angle between the intersection of a fascicle and deep aponeurosis. Fascicle length was defined as the distance from fascicle origin in the deep aponeurosis to insertion in the superficial aponeurosis. The mean value of three images was used to determine pennation angle and fascicle length using the “Angle” tool (Scanlon, Fragala et al. 2014) and “Straight” tool (Erskine, Jones et al. 2009), respectively, of the ImageJ software (1.50b). The typical error and the coefficient of variation for pennation angle and fascicle length were 1.95° (8.74%) and 0.35 cm (4.87%) respectively. Finaly, to reduce assessment bias, ultrasound assessors were blinded to limb protocol during image acquisition and subsequent analyses (muscle cross-sectional area, pennation angle, and fascicle length). Participants were instructed and repeatedly reminded not to disclose limb allocation.

### Muscle biopsies

Muscle tissue samples were obtained from the vastus lateralis by percutaneous needle biopsy performed by an experienced medical professional. The skin over the biopsy site was cleaned with an antiseptic solution, and local anesthesia was administered subcutaneously (2–3 mL of 1% lidocaine). A small incision was made with a sterile scalpel, and a Bergström needle with manual suction was inserted to a depth of approximately 4 cm to obtain ∼100 mg of muscle tissue. Immediately after collection, samples were carefully cleaned of visible blood and connective tissue and subdivided for downstream analyses. Approximately 70–80 mg of tissue was placed into pre-labeled cryotubes for protein and enzymatic activity analyses. An additional 20–30 mg was embedded in optimal cutting temperature (OCT) compound with fibers oriented perpendicular to the cutting surface and rapidly frozen in isopentane cooled with liquid nitrogen for immunohistochemical analyses. All samples were snap-frozen in liquid nitrogen and stored at −80°C for further processing.

### Immunohistochemical analyses

The OCT-embedded samples were sectioned at 7 µm using a criostato (Leica Biosystems, Buffalo Grove, IL, USA), mounted on slides, and stored at -80°C until batch analysis muscle fiber cross-sectional area (fCSA), myonuclei, and satellite cell (SC) number. All samples from a given participant were processed simultaneously to minimize inter-assay variability. Sections were air-dryed (2 h) and fixed with ice-cold acetone at -20°C (5 min). Sections were incubated with 3% H2O2 (10 min), followed by an autofluorescence-quenching reagent (TrueBlack®, cat. no. 23007; Biotium, Fremont, CA, USA) (1 min) and then blocked with a solution of 5% goat serum, 2.5% horse serum, and 0.1% Triton X-100 (1 h). Subsequently, endogenous biotin and streptavidin binding sites were blocked using a Streptavidin/Biotin Blocking Kit (15 min each) before overnight incubation at 4°C with a primary antibody cocktail containing: mouse anti-Pax7 IgG1 (1:20; Supernatant; Developmental Studies Hybridoma Bank; Iowa City, IA, USA) + rabbit anti-dystrophin (1:100; cat. no. GTX57970; GeneTex, Irvine, CA, USA) + mouse anti-MHC I (1:100; BA-D5; RRID: AB_2235587; Developmental Studies Hybridoma Bank, Iowa City, IA, USA) in 2.5% horse serum/PBS. Notably, the same staining protocol has been used prior by our laboratory and is available in an open access journal article elsewhere (Michel, Godwin et al. 2025). The following day, sections were rinsed and incubated with biotin-SP-conjugated goat anti-mouse IgG1 (1:1000; Jackson ImmunoResearch; West Grove, PA, USA) (90 min), followed by incubation in a secondary cocktail containing streptavidin-HRP (SA-HRP) (1:500; cat. no. S-911; Thermo Fisher Scientific) + goat anti-rabbit Alexa Fluor 488 (1:250; Vector Laboratories; Burlingame, CA, USA) + goat anti-rabbit Alexa Fluor 647 (1:250; cat. no. A-21242; Thermo Fisher Scientific) (1 h). Tyramide Signal Amplification (TSA) Alexa Fluor 594 (1:200; cat. no. B-40957; Thermo Fisher Scientific) was applied (20 min), followed by DAPI (1:10,000) (10 min). Finally, slides were mounted in PBS/glycerol (1:1) and digital images (10× and 20× objectives) were captured (Zeiss Axio Imager.M2). SC analysis (Pax7+/DAPI+), fiber typing, fCSA, and myonuclei were quantified manually and via MyoVision software (Wen, Murach et al. 2018). Only regions free of freezing artifacts were analyzed, and an average of ∼540 fibers per sample were included for quantification. A list of antibodies utilized for experiments are provided in Supplementary Material 1.

### Muscle tissue homogenization and protein extraction

Approximately 20 mg of whole muscle tissue was placed in 1.7 mL microcentrifuge tubes with 300 μL of a general cell lysis buffer (cat. no. 9803; Cell Signaling Technology, Danvers, MA, USA). The samples were homogenized using hard plastic pestles and centrifuged at 500 × g at 4°C (5 min). The supernatants were transferred to a new set of microtubes. Approximately 20 µL of the resulting cell lysates were used to determine total protein concentrations using a commercially available bicinchoninic acid (BCA) protein assay kit (cat. no. 23227; Thermo Fisher Scientific) and a spectrophotometer (BioTek Synergy H1, Winooski, VT, USA), following the manufacturer’s instructions.

### Western blotting

The homogenized lysates were added to 4× Laemmli buffer and deionized water (diH2O) at a concentration of 1 µg/µL. The solutions were then denatured at 100°C (5 min) before being stored at -80°C until analysis. Next, the prepared samples (15 μL) were loaded onto 4–15% SDS-polyacrylamide gels (cat. no. 5671085, Criterion TGX; Bio-Rad Laboratories, Hercules, CA, USA) and subjected to electrophoresis at 180 V (50 min) with pre-made 1× SDS-PAGE running buffer (VWR-Avantor, Radnor, PA, USA). The proteins were transferred to pre-activated polyvinylidene difluoride membranes (cat. no. 1620177; Bio-Rad Laboratories) at 200 mA (2 h). The membranes were stained with Ponceau S (10 min) and digitally photographed using a gel documentation system (ChemiDoc Touch; Bio-Rad Laboratories). The membranes were then reactivated in methanol and blocked with 5% skimmed milk powder in Tris-buffered saline with 0.1% Tween-20 (TBST) at room temperature (1 h). The membranes were incubated overnight at 4°C with the following antibodies at a dilution of 1:1000 in TBST with 5% bovine serum albumin (BSA): rabbit anti-MyHC (cat. no: 64038, Cell Signaling Technology); rabbit anti-polyubiquitin (cat. no: 3933, Cell Signaling Technology); rabbit 20S antibody cocktail (cat. no: PW8155, Enzo Life Sciences); rabbit anti-calpain-1 (cat. no: 2556, Cell Signaling Technology); rabbit anti-calpain-2 (cat. no: 70655, Cell Signaling Technology); rabbit anti-LC3 (cat. no: 2775, Cell Signaling Technology); rabbit anti-FOXO1 (cat. no: 9454, Cell Signaling Technology); rabbit anti-FOXO3 (cat. no: 24975, Cell Signaling Technology); rabbit anti-RPS6 (cat. no: 2217, Cell Signaling Technology); rabbit anti-4EBP1 (cat. no: 9644, Cell Signaling Technology); rabbit anti-phospho-4EBP1 (cat. no: 2855, Cell Signaling Technology); rabbit anti-p62 (cat. no: 5114, Cell Signaling Technology); mouse anti-SKIV2L2 (cat. no: sc-515828, Santa Cruz Technology); mouse anti-G3BP1 (cat. no: sc-365338, Santa Cruz Technology); rabbit anti-p70S6K (cat. no: 9234, Cell Signaling Technology); rabbit anti-phospho-p70S6K (cat. no: 2983, Cell Signaling Technology); rabbit anti-mTOR (cat. no: 5536, Cell Signaling Technology); rabbit anti-phospho-mTOR (cat. no: 2971, Cell Signaling Technology). The following day, the membranes were incubated with anti-mouse or anti-rabbit IgG conjugated to horseradish peroxidase (cat. nos.: 7076 and 7074; Cell Signaling Technology) (1 h). The membranes were then developed with a chemiluminescent substrate (Immobilon Forte, cat. no. WBLUF0500; MilliporeSigma, Burlington, MA, USA) protein bands were visualized using a digital imaging system (ChemiDoc Touch, Bio-Rad Laboratories). Representative immunoblots are provided in Supplementary Material 2. Band densities were quantified using Image Lab software (Bio-Rad). Target protein signals were normalized to total protein staining (Ponceau), and for phosphorylated proteins, phospho-to-total protein ratios were calculated.

### Calpain and proteasome activity assays

Calpain and proteasome enzymatic activities were quantified using commercially available luminescence-based assay kits (cat. nos. G8502 and G8622; Promega Corporation, Madison, WI, USA) according to the manufacturer’s instructions. Briefly, aliquots of muscle lysate were diluted 1:10 with diH_2_O and 25 μL were added to white 96-well plates containing the kit-specific reagent mixture. After a 10-min incubation at room temperature, luminescence was measured using a microplate luminometer (BioTek Synergy H1, Winooski, VT, USA). Signals were normalized to total soluble protein concentration determined by BCA assay and expressed as relative luminescence units (RLU) per µg of protein. All samples were analyzed in duplicate. The coefficients of variation between duplicates averaged 2.4% for calpain activity and 3.1% for proteasome activity.

### Real-time quantitative PCR

Total RNA was isolated from frozen vastus lateralis muscle samples (∼20 mg) using TRIzol reagent (Thermo Fisher Scientific, Waltham, MA, USA). Samples were manually homogenized with plastic pestles, and RNA was precipitated using an isopropanol-glycogen mixture. The resulting RNA pellets were washed twice with 75% ethanol, dried, and reconstituted in 30 µL of RNase-free water. RNA concentration and purity were determined in duplicate by spectrophotometry (NanoDrop Lite, Thermo Fisher Scientific). Complementary DNA (cDNA) was synthesized from 2 µg of total RNA using a commercial reverse transcription kit (qScript cDNA SuperMix; Quanta Biosciences, Gaithersburg, MD, USA) according to the manufacturer’s protocol.

Quantitative PCR was performed using SYBR Green chemistry (Quanta Biosciences) on a real-time PCR detection system (Bio-Rad Laboratories, Hercules, CA, USA). Reactions were carried out in a final volume of 20 µL containing gene-specific forward and reverse primers (final concentration: 2 µM each) and 25 ng of cDNA. All samples were analyzed in duplicate. Primer sequences are provided in Supplementary Material 1. Relative gene expression was calculated using the ΔΔCt (Livak and Schmittgen 2001), with glyceraldehyde-3-phosphate dehydrogenase (GAPDH) used as the reference gene. Results are expressed as fold change using the 2^−ΔΔCt^ method.

### Statistical analysis

All statistical analyses were performed using SAS software (version 9.4; SAS Institute Inc., Cary, NC, USA). Visual inspection of residual plots was also performed to verify model assumptions. Primary and secondary outcomes were analyzed using linear mixed-effects models to account for the within-subject unilateral design. Training protocol (VOL20 vs. VOL120), time (pre-intervention, post-intervention [chronic], 0 h and 24 h post-exercise [acute], when applicable), and their interaction were included as fixed effects, and participant was included as a random effect. Model fit was evaluated by inspection of residual distributions and influence diagnostics. Separate models were constructed for each dependent variable, including mCSA, fCSA (mixed, type I and II), myonuclear number, satellite cell content, protein expression/phosphorylation markers, proteolytic activity measures, and gene expression outcomes. Additionally, whenever different between-protocols baseline values were detected, the absolute changes for the dependent variables were compared using an analysis of covariance (ANCOVA). When a significant main effect or interaction was detected, pairwise comparisons were performed using Tukey-adjusted post hoc tests. An unpaired t test was used to compare the number of weekly sets and volume load (sets × repetitions × load [kg]) accumulated throughout the intervention between the experimental protocols. Descriptive data are presented as mean ± standard deviation (SD). Statistical significance was set at *p* < 0.05.

## RESULTS

### Participant flow and adherence

Participant characteristics by protocol are presented in Table 1. Of 138 volunteers screened for eligibility, 29 met the inclusion criteria and consented to participate, and their limbs were randomized to the two training protocols. Of these, 28 participants initiated the training intervention after baseline biopsies, and three withdrew during follow-up due to persistent muscle or joint pain. Thus, 25 participants (18 men and 7 women) completed the intervention and were included in the final analyses. Across the 8-week intervention, a total of 400 supervised training sessions were scheduled, of which 394 were completed, corresponding to a mean adherence rate of 98.5%. A CONSORT flow diagram is shown in figure 2.

**Figure 2.**
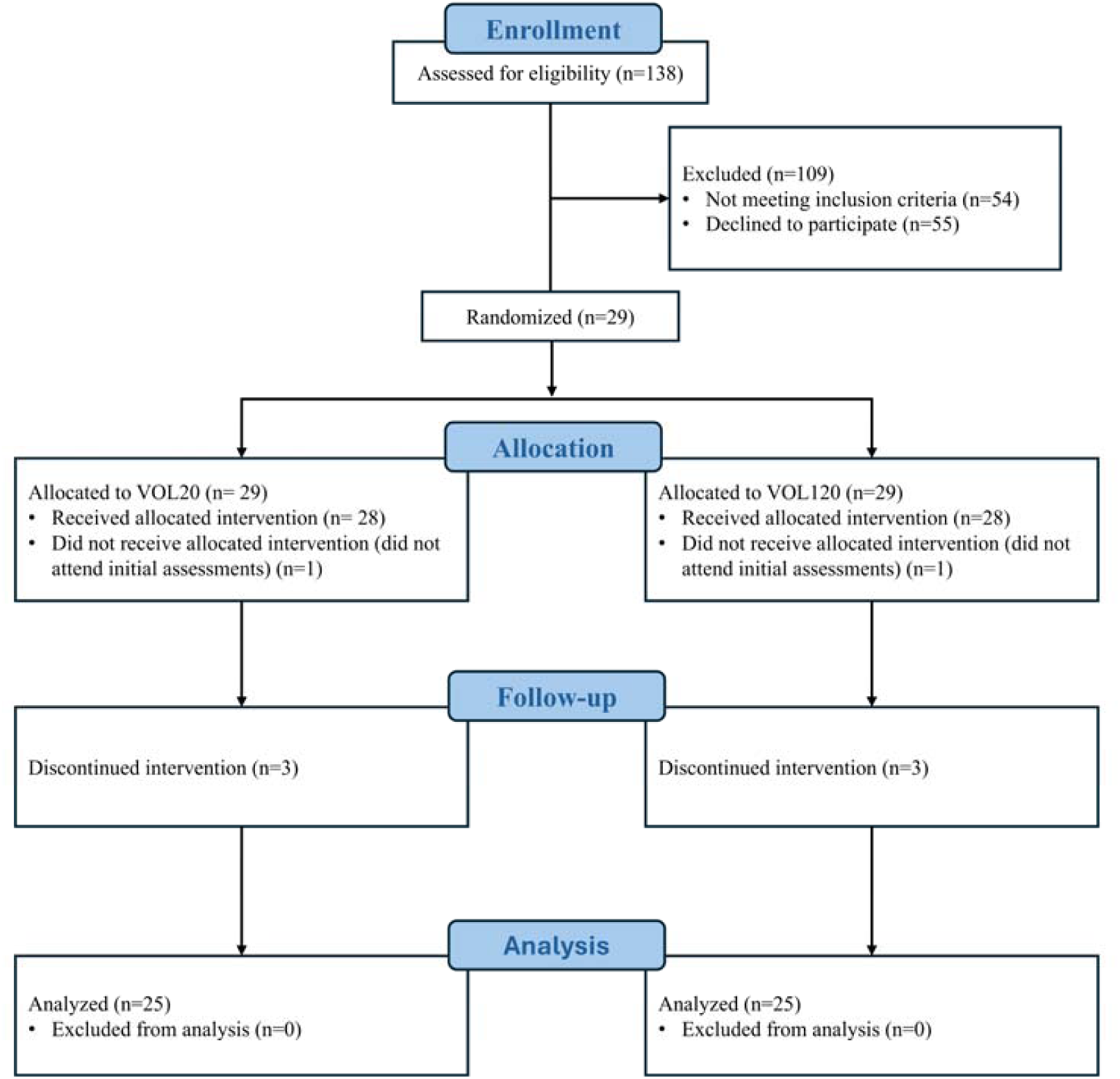
CONSORT flow diagram of participants through the study.

**Table 1.** Baseline sample characteristics.

| Variable | Mean ± Standard deviation |
| --- | --- |
| Age (years) | 25.4 (5.2) |
| Body mass (kg) | 79.3 (19.2) |
| Height (m) | 1.74 (0.11) |
| Training experience (years) | 4.3 (3.6) |
| Number of exercises for quadriceps per week | 4.2 (0.7) |
| Set volume for quadriceps per week | 14.9 (2.6) |
| Legend: kg: kilograms; m: meters. |  |

### Number of sets and volume load

The unpaired t-test revealed a significantly higher number of weekly sets for protocol VOL120 (32.8 ± 5.9) compared to protocol VOL20 (18.0 ± 3.3 sets; *p* < 0.001). The accumulated volume load throughout the intervention was significantly greater in VOL120 (237,125 ± 91,078 kg, *p* < 0.0001) than in VOL20 (138,416 ± 52,6931 kg).

### Dietary adherence

Macronutrient distribution did not significantly change throughout the study period. Mean dietary intake corresponded to 21.8 ± 5.7% of total energy from protein, 49.8 ± 9.5% from carbohydrates, and 28.4 ± 18.1% from fat (detailed data not shown).

### Muscle cross-sectional area, fascicle length and pennation angle

Mixed model analysis indicated a significant main effect of time was observed for vastus lateralis mCSA (*F*_[1,72]_ = 61.02, *p* < 0.001). Both protocols showed significant increases from pre- to post-training (Figure 3A). Additionally, a significant main effect of protocol was observed (*F*_[1,72]_ = 9.71, *p* = 0.002). The VOL120 protocol showed higher mCSA value compared to the VOL20 protocol. No protocol vs. time interaction was observed for mCSA (*F*_[1,72]_ = 1.70, *p* = 0.197). Additionally, no significant main effects or interactions were observed for fascicle length or pennation angle (all *p* > 0.05) (Figure 3B and 3C).

**Figure 3.**
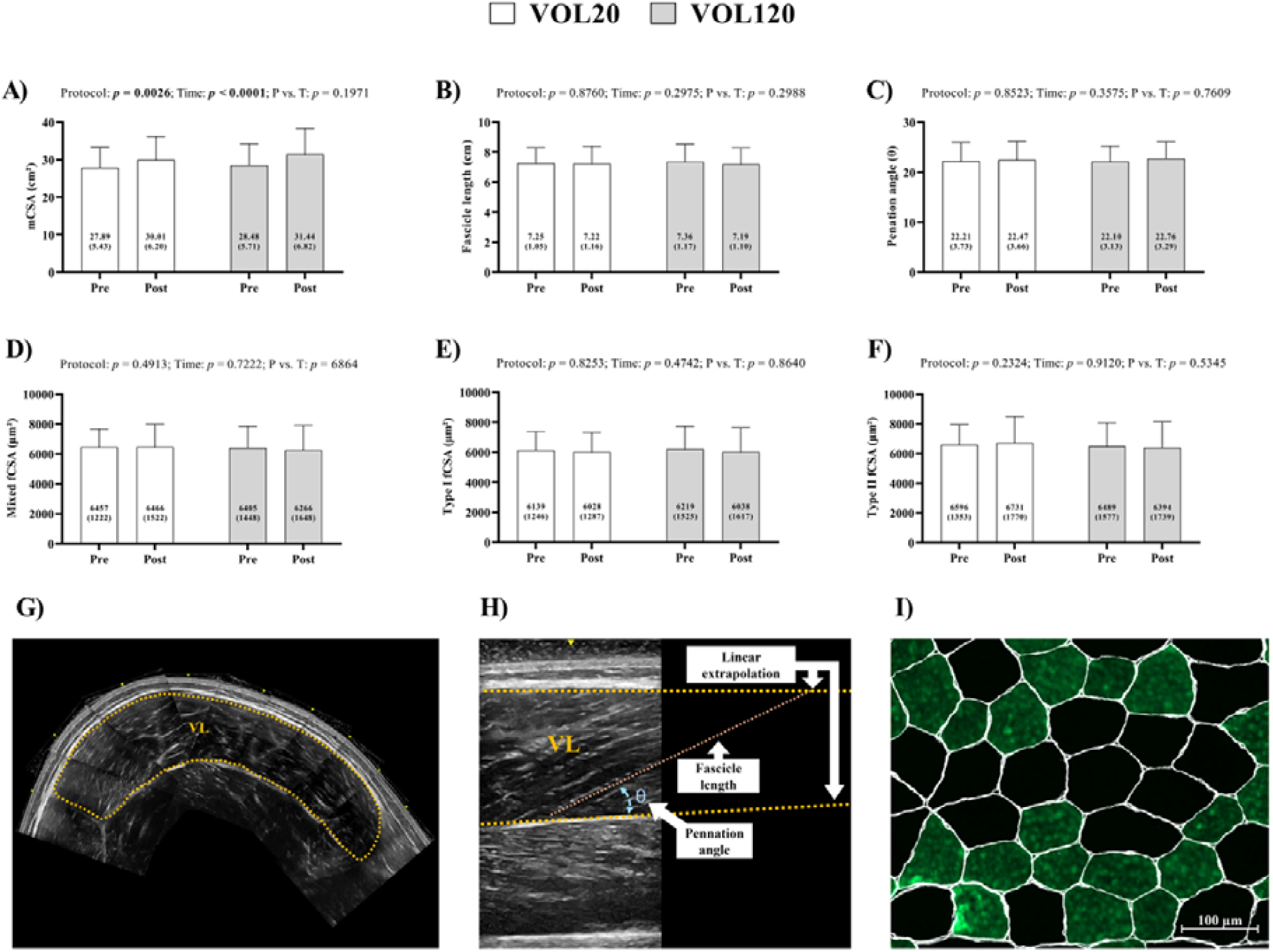
Muscle cross-sectional area (mCSA), fascicle length, pennation angle, and fiber-type-specific morphology for the VOL20 (white bars) and VOL120 (gray bars) protocols before (Pre) and after (Post) 8 weeks of resistance training. (A) mCSA; (B) fascicle length; (C) pennation angle; (D) mixed, (E) type I, and (f) type II fiber cross-sectional areas (fCSA). The lower panels show representative images of the techniques utilized: (G) ultrasound scan of the vastus lateralis used for mCSA measurement; (H) B-mode ultrasound image demonstrating the assessment of pennation angle and fascicle length; and (I) representative immunohistochemical micrograph of type I (green) and type II (black) fibers delineated by dystrophin (white). Scale bar = 100 μm. Values are presented as means (standard deviation).

### Muscle fiber cross-sectional area

Mixed model analysis indicated no significant effects of time, protocol, or protocol vs. time interaction were detected for mixed, type I and type II fCSA (all *p* > 0.05) (Figure 3D, 3E and 3F).

### Satellite cells and myonuclei

Mixed model analysis showed no significant effects of time, protocol, or protocol vs. time interaction were observed for myonuclear content, or satellite cell content for either protocol in chronic or acute analyses (all *p* > 0.05) (Figure 4A-F).

**Figure 4.**
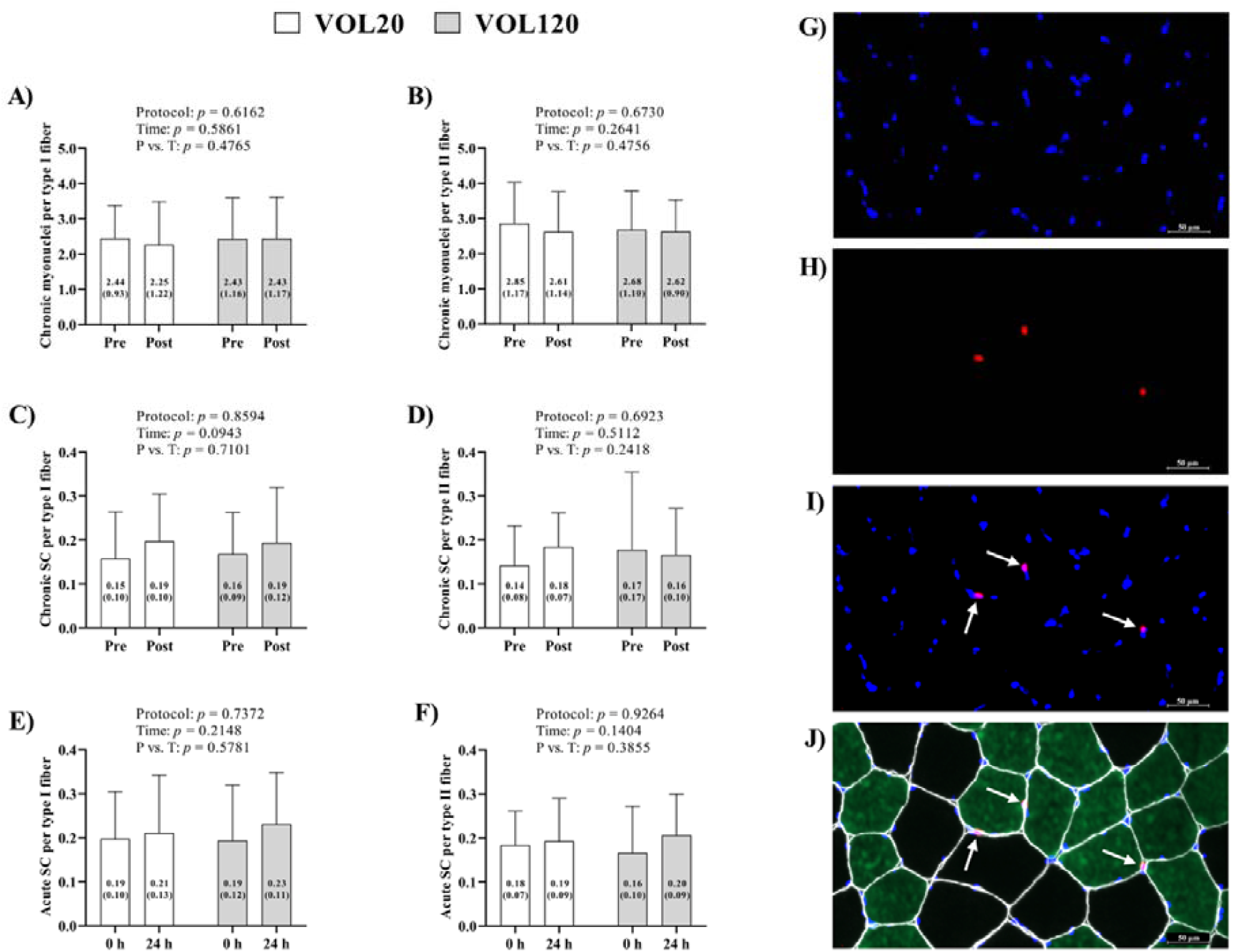
Myonuclear and satellite cell (SC) content per fiber type for the VOL20 (white bars) and VOL120 (gray bars) protocols. For chronic measures (A–D), Pre and Post refer to baseline and 8 weeks of training; for acute measures (E–F), 0 h and 24 h refer to before and after a single session. (A) chronic myonuclei changes per type I fiber; (B) chronic myonuclei changes per type II fiber; (C) chronic satellite cells changes per type I fiber; (D) chronic SC per type II fiber; (E) acute SC per type I fiber; and (F) acute SC per type II fiber. The right panels display representative immunohistochemical micrographs: (G) DAPI staining for nuclei (blue); (H) Pax7 staining for SCs (red); (I) merged DAPI and Pax7 channels, where white arrows indicate SCs (Pax7+/DAPI+); and (J) full composite image including dystrophin (white) and type I fiber (green). Scale bar = 50 μm. Values are presented as means (standard deviation).

### Chronic responses of molecular markers

The mixed-model analysis revealed a significant main effect of time only for CALP-1 (*F*_[1,69]_ = 9.17, *p* = 0.003), CALP-2 (*F*_[1,72]_ = 17.10, *p* < 0.0001), LC3-II (*F*_[1,63]_ = 8.95, *p* = 0.003), and SKIVL2 (*F*_[1,72]_ = 4.46, *p* = 0.038) protein expression, with higher values at post compared with pre-intervention. In contrast, FOXO3 protein expression (*F*_[1,66]_ = 5.00, *p* = 0.028) was higher at pre- compared with post-intervention. A significant main effect of protocol was observed only for CALP-2 (*F*_[1,72]_ = 4.82, *p* = 0.031), with higher protein expression in VOL120 compared with VOL20. Additionally, ANCOVA adjusted for baseline values also indicated a significant between-protocol difference for G3BP1, with higher adjusted values in VOL20 (*p* = 0.005). No significant main effects of protocols and time, or interactions protocol vs. time were detected for any other proteins (all *p* < 0.05). Mean ± SD values for chronic protein expression are presented in Table 2. The mixed-model analysis revealed a significant main effect of time for 45S pre-rRNA gene expression (*F*_[1,69]_ = 4.41, *p* = 0.039), with higher values at post compared with pre-intervention. MSTN mRNA expression also showed a significant main effect of time (*F*_[1,23]_ = 6.98, *p* = 0.014), with lower values at post compared with pre-intervention. A significant main effect of protocol was observed only for 45S pre-rRNA gene expression (*F*_[1,69]_ = 6.82, *p* = 0.011), with higher values in VOL120 compared with VOL20. No significant main effects of protocol and time, or protocol vs. time interaction were observed for the remaining gene targets analyzed (all *p* < 0.05). Mean ± SD values for chronic gene expression data are presented in Table 3. The mixed model analysis revealed no significant main effects of protocol or time, and no protocol vs. time interaction for calpain and proteasome chronic activity.

**Table 2.**
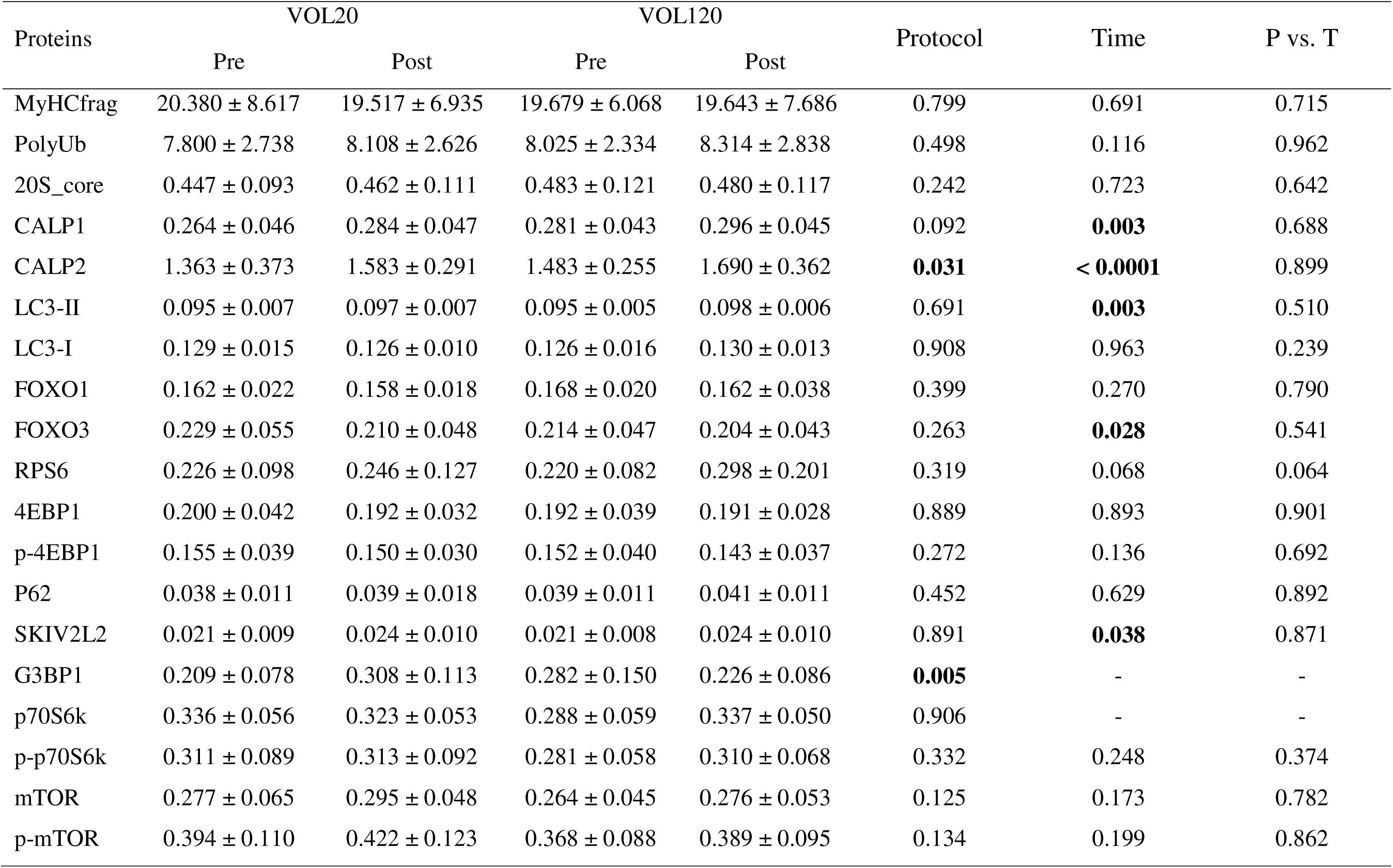
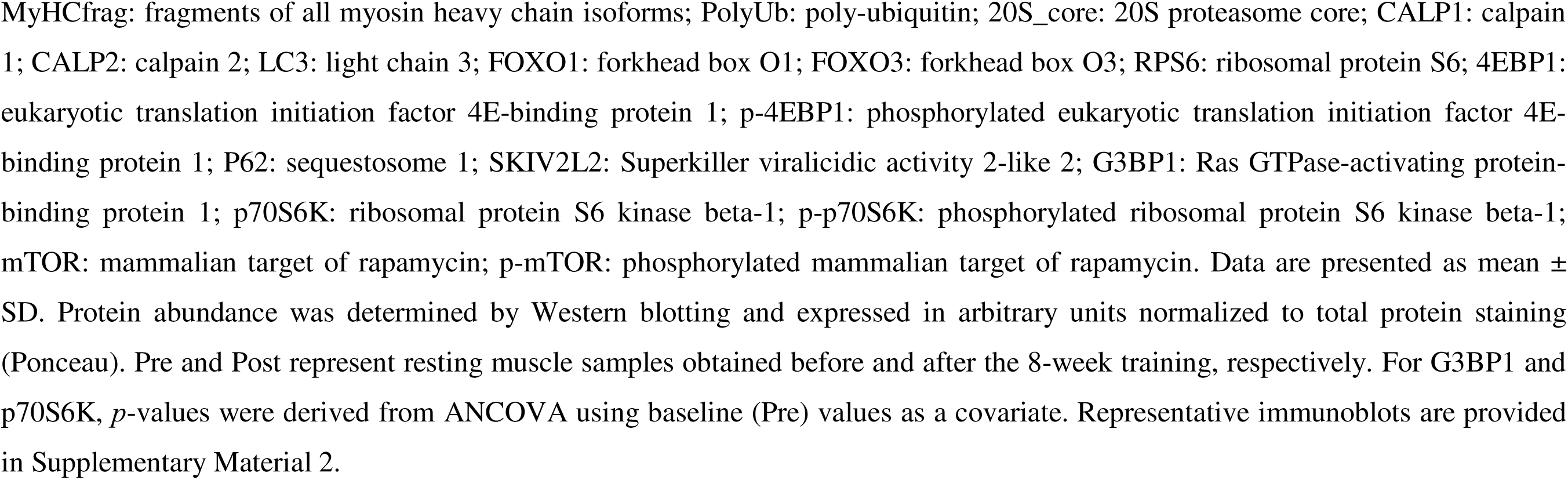
Chronic changes in skeletal muscle protein abundance following modest (+20%; VOL20) versus large (+120%; VOL120) weekly resistance training volume progression.

| Proteins | VOL20 |  | VOL120 |  | Protocol | Time | P vs. T |
| --- | --- | --- | --- | --- | --- | --- | --- |
|  | Pre | Post | Pre | Post |  |  |  |
| MyHCfrag | 20.380 ± 8.617 | 19.517 ± 6.935 | 19.679 ± 6.068 | 19.643 ± 7.686 | 0.799 | 0.691 | 0.715 |
| PolyUb | 7.800 ± 2.738 | 8.108 ± 2.626 | 8.025 ± 2.334 | 8.314 ± 2.838 | 0.498 | 0.116 | 0.962 |
| 20S_core | 0.447 ± 0.093 | 0.462 ± 0.111 | 0.483 ± 0.121 | 0.480 ± 0.117 | 0.242 | 0.723 | 0.642 |
| CALP1 | 0.264 ± 0.046 | 0.284 ± 0.047 | 0.281 ± 0.043 | 0.296 ± 0.045 | 0.092 | **0.003** | 0.688 |
| CALP2 | 1.363 ± 0.373 | 1.583 ± 0.291 | 1.483 ± 0.255 | 1.690 ± 0.362 | **0.031** | **< 0.0001** | 0.899 |
| LC3-II | 0.095 ± 0.007 | 0.097 ± 0.007 | 0.095 ± 0.005 | 0.098 ± 0.006 | 0.691 | **0.003** | 0.510 |
| LC3-I | 0.129 ± 0.015 | 0.126 ± 0.010 | 0.126 ± 0.016 | 0.130 ± 0.013 | 0.908 | 0.963 | 0.239 |
| FOXO1 | 0.162 ± 0.022 | 0.158 ± 0.018 | 0.168 ± 0.020 | 0.162 ± 0.038 | 0.399 | 0.270 | 0.790 |
| FOXO3 | 0.229 ± 0.055 | 0.210 ± 0.048 | 0.214 ± 0.047 | 0.204 ± 0.043 | 0.263 | **0.028** | 0.541 |
| RPS6 | 0.226 ± 0.098 | 0.246 ± 0.127 | 0.220 ± 0.082 | 0.298 ± 0.201 | 0.319 | 0.068 | 0.064 |
| 4EBP1 | 0.200 ± 0.042 | 0.192 ± 0.032 | 0.192 ± 0.039 | 0.191 ± 0.028 | 0.889 | 0.893 | 0.901 |
| p-4EBP1 | 0.155 ± 0.039 | 0.150 ± 0.030 | 0.152 ± 0.040 | 0.143 ± 0.037 | 0.272 | 0.136 | 0.692 |
| P62 | 0.038 ± 0.011 | 0.039 ± 0.018 | 0.039 ± 0.011 | 0.041 ± 0.011 | 0.452 | 0.629 | 0.892 |
| SKIV2L2 | 0.021 ± 0.009 | 0.024 ± 0.010 | 0.021 ± 0.008 | 0.024 ± 0.010 | 0.891 | **0.038** | 0.871 |
| G3BP1 | 0.209 ± 0.078 | 0.308 ± 0.113 | 0.282 ± 0.150 | 0.226 ± 0.086 | **0.005** | - | - |
| p70S6k | 0.336 ± 0.056 | 0.323 ± 0.053 | 0.288 ± 0.059 | 0.337 ± 0.050 | 0.906 | - | - |
| p-p70S6k | 0.311 ± 0.089 | 0.313 ± 0.092 | 0.281 ± 0.058 | 0.310 ± 0.068 | 0.332 | 0.248 | 0.374 |
| mTOR | 0.277 ± 0.065 | 0.295 ± 0.048 | 0.264 ± 0.045 | 0.276 ± 0.053 | 0.125 | 0.173 | 0.782 |
| p-mTOR | 0.394 ± 0.110 | 0.422 ± 0.123 | 0.368 ± 0.088 | 0.389 ± 0.095 | 0.134 | 0.199 | 0.862 |

MyHCfrag: fragments of all myosin heavy chain isoforms; PolyUb: poly-ubiquitin; 20S\_core: 20S proteasome core; CALP1: calpain 1; CALP2: calpain 2; LC3: light chain 3; FOXO1: forkhead box O1; FOXO3: forkhead box O3; RPS6: ribosomal protein S6; 4EBP1: eukaryotic translation initiation factor 4E-binding protein 1; p-4EBP1: phosphorylated eukaryotic translation initiation factor 4E-binding protein 1; P62: sequestosome 1; SKIV2L2: Superkiller viralicidic activity 2-like 2; G3BP1: Ras GTPase-activating protein-binding protein 1; p70S6K: ribosomal protein S6 kinase beta-1; p-p70S6K: phosphorylated ribosomal protein S6 kinase beta-1; mTOR: mammalian target of rapamycin; p-mTOR: phosphorylated mammalian target of rapamycin. Data are presented as mean ± SD. Protein abundance was determined by Western blotting and expressed in arbitrary units normalized to total protein staining (Ponceau). Pre and Post represent resting muscle samples obtained before and after the 8-week training, respectively. For G3BP1 and p70S6K, *p*-values were derived from ANCOVA using baseline (Pre) values as a covariate. Representative immunoblots are provided in Supplementary Material 2.

**Table 3.** Chronic changes in skeletal muscle gene expression following modest (+20%; VOL20) versus large (+120%; VOL120) weekly resistance training volume progression.

| Genes | VOL20 |  | VOL120 |  | Protocol | Time | P vs. T |
| --- | --- | --- | --- | --- | --- | --- | --- |
|  | Pre | Post | Pre | Post |  |  |  |
| 18S | 1.155 ± 0.753 | 0.989 ± 0.383 | 1.220 ± 0.907 | 1.111 ± 0.640 | 0.428 | 0.164 | 0.699 |
| 28S | 1.178 ± 0.607 | 1.173 ± 0.556 | 1.292 ± 0.838 | 1.739 ± 1.274 | 0.058 | 0.223 | 0.210 |
| 45S pre-rRNA | 1.179 ± 0.666 | 1.359 ± 0.837 | 1.448 ± 1.199 | 2.000 ± 1.472 | **0.011** | **0.039** | 0.289 |
| MSTN | 1.312 ± 1.228 | 0.770 ± 0.438 | 1.252 ± 0.894 | 0.793 ± 0.506 | 0.879 | **0.014** | 0.656 |
| IGF-1 | 1.292 ± 0.971 | 1.364 ± 1.162 | 1.421 ± 1.226 | 0.815 ± 0.533 | 0.268 | 0.160 | 0.075 |
| IL-15 | 1.164 ± 0.871 | 1.051 ± 0.520 | 1.123 ± 0.608 | 0.941 ± 0.391 | 0.480 | 0.208 | 0.669 |
| FST | 1.232 ± 0.934 | 2.675 ± 6.712 | 1.428 ± 1.621 | 1.293 ± 1.422 | 0.445 | 0.370 | 0.264 |
| Total RNA | 1297 ± 721 | 1330 ± 216 | 1186 ± 362 | 1346 ± 249 | 0.579 | 0.290 | 0.433 |

18S: 18S ribosomal RNA subunit; 28S: 28S ribosomal RNA subunit; 45S pre-rRNA: 45S pre-ribosomal RNA; MSTN: myostatin; IGF-1: insulin like growth factor 1; IL-15: interleukin 15; FST: follistatin; Total RNA: total RNA concentration. Data are presented as mean ± SD (n = 25). Relative gene expression was determined by real-time quantitative PCR and calculated using the $2^{-\Delta\Delta C_t}$ method with GAPDH as the reference gene. Total RNA was calculated as concentration/weight\*volume loaded (ng/mg tissue). Data are presented as mean ± SD. Pre and Post represent resting muscle samples obtained before and after the 8-week training, respectively.

### Acute responses of molecular markers

The mixed-model analysis revealed significant main effects of time for MyHCfrag (*F* _[1,71]_ = 4.16, *p* = 0.045) and p-p70S6k (*F*_[1,66]_ = 14.36, *p* = 0.0003) protein expression with higher values at 24 h post-exercise compared with 0 h. Significant main effects of time were also observed for p70S6K (*F* _[1,29]_ = 12.61, *p* = 0.001), CALP-1 (*F*_[1,71]_ = 17.06, *p* < 0.0001), CALP-2 (*F*_[1,71]_ = 4.16, *p* = 0.045), SKIV2L2 (*F*_[1,71]_ = 10.07, *p* = 0.002), 4EBP1 (*F*_[1,74]_ = 4.65, *p* = 0.035), p-4EBP1(*F*_[1,73]_ = 15.14, *p* = 0.0002), and (*F*_[1,29]_ = 12.67, *p* =0.001), but with lower protein expression at 24 h post-exercise compared with 0 h. Significant main effects of protocol were only observed for CALP-1 (*F*_[1,70]_ = 5.03, *p* = 0.028), and RPS6 (*F*_[1,24]_ = 5.89, *p* = 0.023), with higher protein expression in VOL120 compared with VOL20. No significant main effects of protocol and time, or interaction protocol vs. time were detected for any other proteins (all *p* < 0.05). Mean ± SD values for acute protein expression are presented in Table 4. The mixed-model analysis revealed a significant main effect of time only for 45S pre-rRNA gene expression (*F*_[1,22]_ = 9.10, *p* = 0.006), 28S rRNA gene expression (*F*_[1,31]_ = 4.82, *p* = 0.035), and FST mRNA expression (*F*_[1,22]_ = 4.54, *p* = 0.044), with higher values at 24 h post-exercise compared with 0 h. No significant main effects of time or protocol, and no protocol vs. time interactions, were observed for the remaining gene targets analyzed. Mean ± SD values for acute gene expression are presented in Table 5. The mixed model analysis revealed a significant main effect of time for calpain (*F*_[1,22]_ = 15.98; *p* = 0.001), and for proteasome activity (*F*_[1,22]_ = 6.57; *p* = 0.017), both with higher values at 24h-post exercise compared with 0h. A significant main effect of protocol was observed only for proteasome activity (*F*_[1,23]_ = 7.76; *p* = 0.010), with higher values for VOL20 compared to VOL120. No protocol vs. time interaction was observed for the acute calpain and proteasome activities.

**Table 4.**
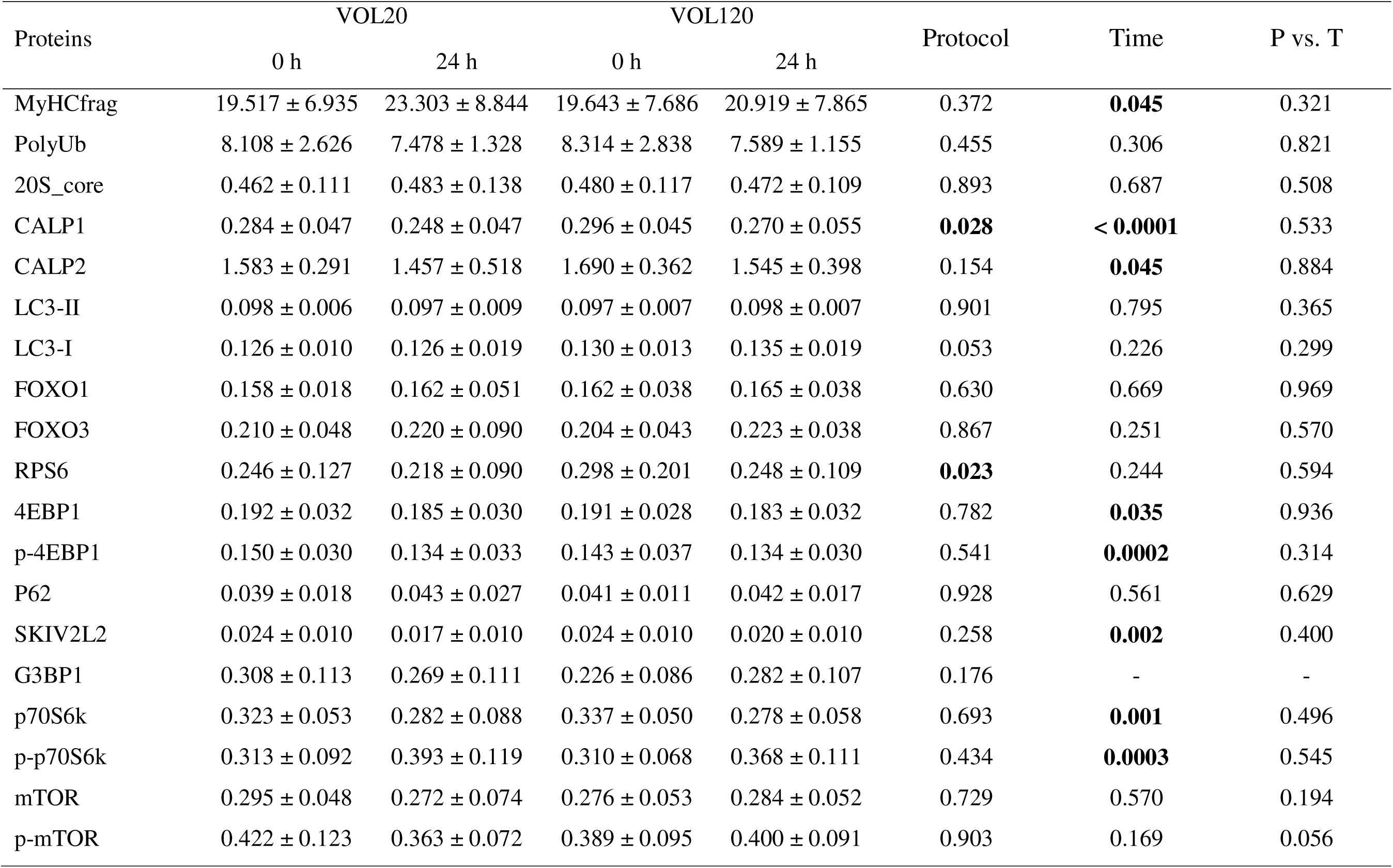
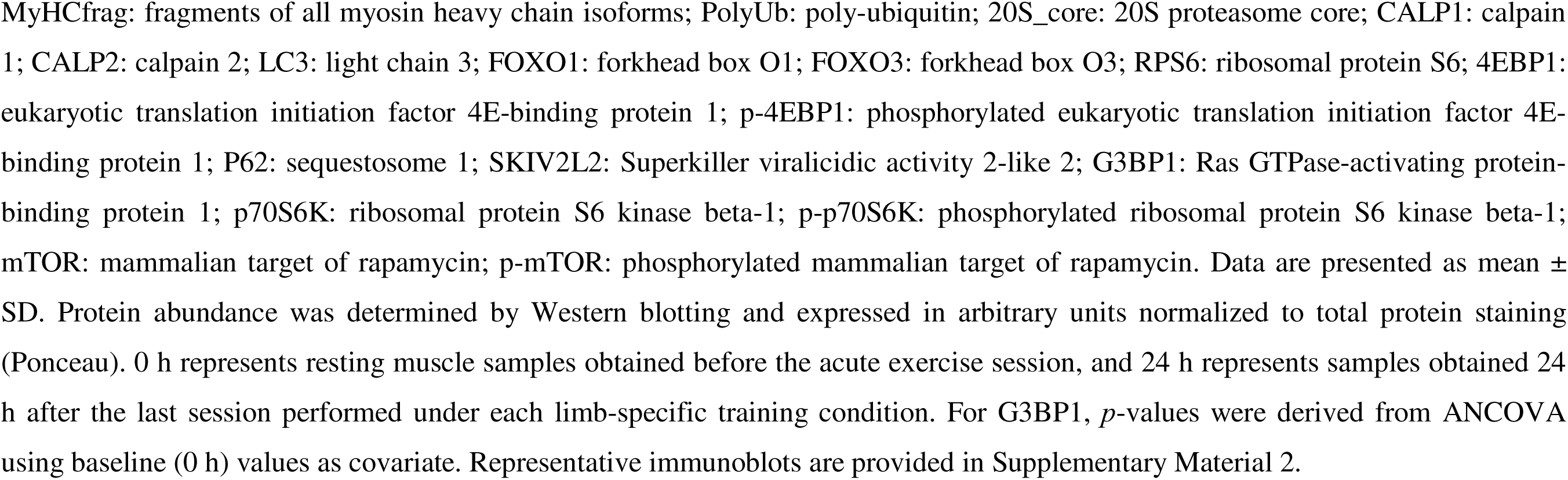
Acute changes in skeletal muscle protein abundance following modest (+20%; VOL20) versus large (+120%; VOL120) weekly resistance training volume progression.

| Proteins | VOL20 |  | VOL120 |  | Protocol | Time | P vs. T |
| --- | --- | --- | --- | --- | --- | --- | --- |
|  | 0 h | 24 h | 0 h | 24 h |  |  |  |
| MyHCfrag | 19.517 ± 6.935 | 23.303 ± 8.844 | 19.643 ± 7.686 | 20.919 ± 7.865 | 0.372 | **0.045** | 0.321 |
| PolyUb | 8.108 ± 2.626 | 7.478 ± 1.328 | 8.314 ± 2.838 | 7.589 ± 1.155 | 0.455 | 0.306 | 0.821 |
| 20S_core | 0.462 ± 0.111 | 0.483 ± 0.138 | 0.480 ± 0.117 | 0.472 ± 0.109 | 0.893 | 0.687 | 0.508 |
| CALP1 | 0.284 ± 0.047 | 0.248 ± 0.047 | 0.296 ± 0.045 | 0.270 ± 0.055 | **0.028** | **< 0.0001** | 0.533 |
| CALP2 | 1.583 ± 0.291 | 1.457 ± 0.518 | 1.690 ± 0.362 | 1.545 ± 0.398 | 0.154 | **0.045** | 0.884 |
| LC3-II | 0.098 ± 0.006 | 0.097 ± 0.009 | 0.097 ± 0.007 | 0.098 ± 0.007 | 0.901 | 0.795 | 0.365 |
| LC3-I | 0.126 ± 0.010 | 0.126 ± 0.019 | 0.130 ± 0.013 | 0.135 ± 0.019 | 0.053 | 0.226 | 0.299 |
| FOXO1 | 0.158 ± 0.018 | 0.162 ± 0.051 | 0.162 ± 0.038 | 0.165 ± 0.038 | 0.630 | 0.669 | 0.969 |
| FOXO3 | 0.210 ± 0.048 | 0.220 ± 0.090 | 0.204 ± 0.043 | 0.223 ± 0.038 | 0.867 | 0.251 | 0.570 |
| RPS6 | 0.246 ± 0.127 | 0.218 ± 0.090 | 0.298 ± 0.201 | 0.248 ± 0.109 | **0.023** | 0.244 | 0.594 |
| 4EBP1 | 0.192 ± 0.032 | 0.185 ± 0.030 | 0.191 ± 0.028 | 0.183 ± 0.032 | 0.782 | **0.035** | 0.936 |
| p-4EBP1 | 0.150 ± 0.030 | 0.134 ± 0.033 | 0.143 ± 0.037 | 0.134 ± 0.030 | 0.541 | **0.0002** | 0.314 |
| P62 | 0.039 ± 0.018 | 0.043 ± 0.027 | 0.041 ± 0.011 | 0.042 ± 0.017 | 0.928 | 0.561 | 0.629 |
| SKIV2L2 | 0.024 ± 0.010 | 0.017 ± 0.010 | 0.024 ± 0.010 | 0.020 ± 0.010 | 0.258 | **0.002** | 0.400 |
| G3BP1 | 0.308 ± 0.113 | 0.269 ± 0.111 | 0.226 ± 0.086 | 0.282 ± 0.107 | 0.176 | - | - |
| p70S6k | 0.323 ± 0.053 | 0.282 ± 0.088 | 0.337 ± 0.050 | 0.278 ± 0.058 | 0.693 | **0.001** | 0.496 |
| p-p70S6k | 0.313 ± 0.092 | 0.393 ± 0.119 | 0.310 ± 0.068 | 0.368 ± 0.111 | 0.434 | **0.0003** | 0.545 |
| mTOR | 0.295 ± 0.048 | 0.272 ± 0.074 | 0.276 ± 0.053 | 0.284 ± 0.052 | 0.729 | 0.570 | 0.194 |
| p-mTOR | 0.422 ± 0.123 | 0.363 ± 0.072 | 0.389 ± 0.095 | 0.400 ± 0.091 | 0.903 | 0.169 | 0.056 |

MyHCfrag: fragments of all myosin heavy chain isoforms; PolyUb: poly-ubiquitin; 20S\_core: 20S proteasome core; CALP1: calpain 1; CALP2: calpain 2; LC3: light chain 3; FOXO1: forkhead box O1; FOXO3: forkhead box O3; RPS6: ribosomal protein S6; 4EBP1: eukaryotic translation initiation factor 4E-binding protein 1; p-4EBP1: phosphorylated eukaryotic translation initiation factor 4E-binding protein 1; P62: sequestosome 1; SKIV2L2: Superkiller viralicidic activity 2-like 2; G3BP1: Ras GTPase-activating protein-binding protein 1; p70S6K: ribosomal protein S6 kinase beta-1; p-p70S6K: phosphorylated ribosomal protein S6 kinase beta-1; mTOR: mammalian target of rapamycin; p-mTOR: phosphorylated mammalian target of rapamycin. Data are presented as mean ± SD. Protein abundance was determined by Western blotting and expressed in arbitrary units normalized to total protein staining (Ponceau). 0 h represents resting muscle samples obtained before the acute exercise session, and 24 h represents samples obtained 24 h after the last session performed under each limb-specific training condition. For G3BP1, *p*-values were derived from ANCOVA using baseline (0 h) values as covariate. Representative immunoblots are provided in Supplementary Material 2.

**Table 5.** Acute changes in skeletal muscle gene expression following modest (+20%; VOL20) versus large (+120%; VOL120) weekly resistance training volume progression.

| Genes | VOL20 |  | VOL120 |  | Protocol | Time | P vs. T |
| --- | --- | --- | --- | --- | --- | --- | --- |
|  | 0 h | 24 h | 0 h | 24 h |  |  |  |
| 18S | 1.075 ± 0.418 | 1.338 ± 0.791 | 1.155 ± 0.663 | 1.201 ± 0.785 | 0.805 | 0.215 | 0.350 |
| 28S | 1.136 ± 0.538 | 1.700 ± 0.928 | 1.255 ± 0.919 | 1.492 ± 0.910 | 0.798 | **0.035** | 0.363 |
| 45S pre-rRNA | 1.252 ± 0.770 | 2.271 ± 1.811 | 1.300 ± 0.955 | 1.911 ± 1.558 | 0.479 | **0.006** | 0.416 |
| MSTN | 1.168 ± 0.662 | 0.860 ± 1.037 | 1.255 ± 0.800 | 1.021 ± 1.146 | 0.476 | 0.112 | 0.831 |
| IGF-1 | 1.262 ± 1.075 | 1.622 ± 1.722 | 1.157 ± 0.756 | 1.557 ± 1.317 | 0.749 | 0.143 | 0.937 |
| IL-15 | 1.122 ± 0.556 | 1.300 ± 0.593 | 1.087 ± 0.451 | 1.236 ± 0.691 | 0.623 | 0.142 | 0.885 |
| FST | 2.357 ± 5.913 | 4.691 ± 7.181 | 1.381 ± 1.518 | 3.771 ± 6.485 | 0.492 | **0.044** | 0.974 |
| Total RNA | 1330 ± 216 | 1281 ± 269 | 1346 ± 249 | 1448 ± 502 | 0.163 | 0.783 | 0.133 |

18S: 18S ribosomal RNA subunit; 28S: 28S ribosomal RNA subunit; 45S pre-rRNA: 45S pre-ribosomal RNA subunit; MSTN: myostatin; IGF-1: insulin like growth factor 1; IL-15: interleukin 15; FST: follistatin; Total RNA: total RNA concentration. Data are presented as mean ± SD (n = 25). Relative gene expression was determined by real-time quantitative PCR and calculated using the $2^{-\Delta\Delta Ct}$ method with GAPDH as the reference gene. Total RNA was calculated as concentration/weight\*volume loaded (ng/mg tissue). Data are presented as mean ± SD. 0 h and 24 h represent resting muscle samples obtained before and after the last session performed under each limb-specific training condition.

## DISCUSSION

We sought to determine whether a large and abrupt increase in weekly RT volume would attenuate hypertrophic adaptations and alter molecular markers associated with anabolic and proteolytic regulation in resistance-trained individuals. Contrary to our initial hypothesis, the main findings indicate that a +120% increase in weekly training volume did not impair skeletal muscle hypertrophy at either the mCSA or fCSA level compared with a modest +20% weekly increase. Both volume-progression strategies produced significant increases in vastus lateralis mCSA, and no protocol × time interaction was detected, indicating a similar temporal hypertrophic response between protocols despite differences in absolute training volume. Moreover, most acute and chronic molecular markers related to anabolic signaling, proteolytic pathways, autophagy-related signaling, as well as ribosome biogenesis and turnover exhibited comparable responses between protocols. Collectively, these findings indicate that, under the present experimental protocols, resistance-trained individuals retain an adaptive capacity following large abrupt increases in weekly training volume, without evidence of impaired morphological changes or between-protocols differences in the molecular markers assessed.

Evidence in resistance-trained individuals does not consistently indicate a hypertrophic advantage with higher volumes. While some independent studies report greater gains with higher volumes (Schoenfeld, Ogborn et al. 2017, Schoenfeld, Contreras et al. 2019, Brigatto, Lima et al. 2022), others show comparable adaptations despite substantial differences in prescribed volume (Heaselgrave, Blacker et al. 2019, Aube, Wadhi et al. 2022, Enes, EO et al. 2024, Moreno, Sampson et al. 2024). In line with the latter, Aube et al. (2022) observed similar vastus lateralis muscle thickness increases despite markedly different relative increases in weekly sets (∼30% vs. ∼98% above prior training volume), although progression magnitude was not experimentally imposed, which limits causal inference. Similarly, Barsuhn et al. (2025) and Enes et al. (2024) also reported comparable hypertrophic responses across different volume progressions in trained individuals. Collectively, these findings indicate that the magnitude and direction of any volume-related effect in trained muscle is contingent on experimental design and biological context rather than uniformly expressed across studies (Camargo, Bittencourt et al. 2025).

Several methodological factors likely contribute to such discrepant findings. First, in many prior investigations, training volume was prescribed in absolute terms without reference to participants’ habitual volume, such that the effective relative overload likely varied substantially between individuals (Schoenfeld, Contreras et al. 2019, Brigatto, Lima et al. 2022). In the present study, both interventions were anchored to each participant’s prior weekly set volume, allowing the comparison to more directly examine the magnitude of relative progression (+20% vs +120%). While this feature does not by itself explain divergent literature outcomes, it reduces an important source of prescription heterogeneity that is often unaccounted for in volume-comparison studies (Scarpelli, Nobrega et al. 2020). Second, most prior investigations have employed between-subject designs, which are inherently more susceptible to inter-individual variability in hypertrophic responsiveness driven by genetic, molecular, and training-history factors (Radaelli, Fleck et al. 2015, Schoenfeld, Contreras et al. 2019, Brigatto, Lima et al. 2022, Enes, EO et al. 2024). The unilateral within-subject design adopted here reduces biological heterogeneity and increases sensitivity for detecting true protocol-dependent effects (MacInnis, McGlory et al. 2017). Under these more controlled conditions, the absence of a protocol × time interaction suggests that the magnitude of mCSA increase from pre- to post-intervention did not differ detectably between progression magnitudes. Importantly, although a main effect protocol was detected for mCSA, the between-protocol difference in mCSA increases (2.7%) was small and close to the measurement coefficient of variation based on the typical error of the ultrasound method (2.3%). This was accompanied by a small standardized effect favoring VOL120, but with a wide 95% CI crossing zero (ES = −0.36, 95% CI −1.35 to 0.64; based on VOL120 − VOL20 pre-to-post percent changes in mCSA), reinforcing a cautious interpretation of protocol superiority. Moreover, hypertrophy was not accompanied by detectable changes in fascicle length or pennation angle, in agreement with prior findings in resistance-trained cohorts undergoing moderate-duration RT interventions (Mangine, Redd et al. 2018, Vann, Sexton et al. 2022). At the muscle fiber level, neither protocol produced significant increases in fCSA. While this may appear counterintuitive given the mCSA increases, similar dissociations between macroscopic and microscopic hypertrophy measures have been previously reported in trained individuals (Bjornsen, Wernbom et al. 2019, Carroll, Bazyler et al. 2019, Haun, Vann et al. 2019). Fiber-level adaptations assessed via immunohistochemistry typically show greater sampling variability and lower sensitivity to detect change than imaging-based whole-muscle measures, particularly in previously trained muscle (Haun, Vann et al. 2019). Taken together, these observations indicate that hypertrophic responses to higher RT volumes are context-dependent and highlight the importance of examining underlying cellular and molecular regulatory pathways when interpreting volume-related adaptations.

The regulation of skeletal muscle hypertrophy in response to different resistance training volumes depends on the coordinated interaction between anabolic and catabolic molecular pathways that collectively determine net protein balance and tissue remodeling (Roberts, McCarthy et al. 2023). Experimental evidence indicates that increases in exercise volume can augment anabolic signaling, including p70S6K phosphorylation (Ahtiainen, Walker et al. 2015), satellite cell activation (Hanssen, Kvamme et al. 2013), and ribosome biogenesis (Hammarstrom, Ofsteng et al. 2020). Yet other studies suggest that the muscle protein synthetic response reaches a ceiling beyond which further increases in volume do not proportionally enhance anabolic signaling (Kumar, Atherton et al. 2012). In parallel, very high training volumes have been proposed to increase activation of proteolytic systems, including markers of the ubiquitin–proteasome and autophagy-related pathways, potentially shifting protein balance away from net accretion (Takegaki, Ogasawara et al. 2017, Takegaki, Ogasawara et al. 2019). In this context, the present study was designed to integrate molecular and morphological assessments to examine whether large abrupt increases in weekly training volume disrupt regulatory signaling in trained muscle. Importantly, the acute molecular trial was performed only after completion of the training intervention to minimize the confounding influence of early-phase exercise-induced muscle damage and remodeling, which are known to dissociate acute molecular signaling from hypertrophic outcomes during the initial weeks of RT (Damas, Libardi et al. 2018, Damas, Angleri et al. 2019, Angleri, Damas et al. 2022). Overall, our data indicate that most anabolic and catabolic signaling markers responded similarly under modest and large volume progressions, consistent with the comparable hypertrophic adaptations observed between protocols.

With respect to chronic anabolic signaling, most protein and gene targets associated with mTORC1 signaling and translational regulation were not significantly altered by the intervention, nor did they differ between protocols. Although much of the literature on chronic molecular responses derives from untrained samples, our findings align with prior reports in resistance-trained cohorts showing limited sustained changes in canonical anabolic signaling proteins following high-volume training phases (Haun, Vann et al. 2019). Satellite cell and myonuclear content were likewise unchanged, in agreement with previous studies in trained individuals undergoing RT interventions (Haun, Vann et al. 2019, Angleri, Damas et al. 2022). One plausible interpretation is that previously trained muscle may already possess an expanded myonuclear domain from prior training exposure, reducing the necessity for additional myonuclear accretion over moderate-duration interventions (Petrella, Kim et al. 2008, Conceicao, Vechin et al. 2018). Among the markers related to ribosome biogenesis and RNA homeostasis, 45S pre-rRNA and SKIV2L2 protein levels were the only targets upregulated following the 8-week intervention period. The first and rate-limiting step of ribosome biogenesis is the transcription of rDNA into the 47S pre-rRNA transcript, which is subsequently processed to 45S pre-rRNA and eventually into the mature 28S, 18S, and 5.8S rRNAs (Figueiredo, Caldow et al. 2015). The increase in basal (i.e., 72-hour post bout) 45S pre-rRNA expression observed here is consistent with previous rodent mechanical overload (von Walden, Casagrande et al. 2012) and human training (Figueiredo, Caldow et al. 2015) studies, suggesting that mechanical overload can chronically stimulate increases in translational capacity. In parallel, SKIV2L2 is an RNA helicase and a core cofactor of the nuclear RNA exosome involved in RNA surveillance and rRNA processing/turnover (Lingaraju, Johnsen et al. 2019). Its upregulation may therefore reflect modulation of RNA quality-control dynamics rather than increased ribosome biogenesis per se. Along similar lines, G3BP1, an RNA-binding protein central to stress granule assembly and RNA homeostasis (Panas, Ivanov et al. 2016), displayed a protocol effect, although this was not accompanied by divergent hypertrophic outcomes. Collectively, these results suggest subtle modulation of RNA regulatory pathways with sustained loading, even when morphological and canonical anabolic signaling outcomes are comparable between volume progressions. Acute anabolic signaling was not enhanced under the higher-volume protocol. At 24 h post-exercise, most anabolic signaling targets were similar between conditions, and p70S6K protein abundance was reduced despite a time-dependent increase in p-p70S6K. This contrasts with studies reporting volume-dependent increases in anabolic signaling, after higher per-session set numbers (Burd, Holwerda et al. 2010, Terzis, Spengos et al. 2010, Kumar, Atherton et al. 2012, Ahtiainen, Walker et al. 2015, Hammarstrom, Ofsteng et al. 2020). However, much of this literature involves untrained participants and assessments conducted within the early post-exercise window. Notwithstanding, our findings extend prior work by indicating that, in trained individuals, large abrupt increases in weekly volume do not necessarily amplify chronic and acute anabolic signaling responses, which is consistent with the similar hypertrophic outcomes observed between protocols.

For the chronic proteolytic and remodeling-related signaling, the molecular profile did not indicate a maladaptive catabolic shift under the higher-volume progression. MSTN gene expression decreased after the intervention, independent of protocol, in agreement with evidence linking reduced myostatin-related markers with RT–induced hypertrophy (Roth, Martel et al. 2003, Kim, Petrella et al. 2007), although this response in trained individuals have been inconsistent (Santos, Lamas et al. 2015). Calpain-1 and calpain-2 protein content increased chronically, whereas enzymatic calpain activity did not, a dissociation that is physiologically plausible given the tight regulation of calpain activity by Ca² availability, autolysis, subcellular localization, and calpastatin inhibition (Delaney, Miller et al. 2014, Michel, Godwin et al. 2023). With respect to autophagy-related markers, LC3-I was unchanged, whereas LC3-II protein levels were chronically increased. This apparent divergence between isoforms has been reported previously (Brandt, Dethlefsen et al. 2017) and may reflect increased LC3-I production with subsequent conversion to LC3-II, rather than a simple accumulation pattern. Members of the FOXO family are well-established transcription factors involved in whole-body energy metabolism, skeletal muscle mass regulation, and substrate utilization (Lundell, Massart et al. 2019). In the present study, FOXO3 protein levels were chronically reduced, whereas FOXO1 remained unchanged. This differential behavior is mechanistically plausible, as FOXO3 is highly responsive to cellular stress and is strongly associated with autophagy-related gene regulation, whereas FOXO1 has been more closely linked to overt catabolic conditions such as prolonged disuse or severe energy deprivation (Lundell, Massart et al. 2019). Accordingly, the absence of change in FOXO1 supports the interpretation that the training intervention did not induce a chronic catabolic state, while the reduction in FOXO3 likely reflects adaptive remodeling rather than disproportionate proteolytic activation. This interpretation is further supported by the largely unchanged levels of polyubiquitinated proteins and proteasome subunits after the intervention, in agreement with prior studies in similar trained cohorts (Haun, Vann et al. 2019, Vann, Sexton et al. 2022). Consistent with the chronic findings, acute markers associated with ubiquitin–proteasome and autophagy-related pathways were largely unchanged across protocols and time points. This agrees with several prior studies reporting minimal acute modulation of these pathways following resistance exercise in humans (Delaney, Miller et al. 2014, Michel, Godwin et al. 2023). Notably, the only prior experimental study directly comparing proteolytic signaling across distinct RT volumes was conducted in rodents, and no volume effect on ubiquitin or LC3 responses were reported (Ogasawara, Arihara et al. 2017). Taken together, the present acute and chronic proteolytic signaling data indicate that a large weekly volume progression does not disproportionally alter markers associated with protein breakdown pathways, supporting the interpretation that hypertrophic adaptations were not constrained by a volume-induced catabolic imbalance under the protocols studied.

### Experimental considerations

The present study is not without limitations. Although dietary habits were recorded prior to each muscle biopsy session, daily macronutrient intake was not strictly controlled throughout the intervention. Nutritional variability can influence hypertrophic outcomes. However, available evidence suggests that inter-individual biological factors and training-related variables contribute more strongly to RT–induced hypertrophy than self-reported macronutrient intakes (Thalacker-Mercer, Petrella et al. 2009, Mobley, Haun et al. 2018, Angleri, Damas et al. 2022). Importantly, the within-subject design reduces the likelihood that nutritional variability differentially biased the comparison between volume protocols, as both protocols were implemented within the same individuals. In addition, participants were instructed to maintain their habitual dietary patterns, and a standardized 30 g whey protein supplement was provided after each training session to support post-exercise anabolic responses (Witard, Jackman et al. 2014). A second limitation relates to participant characteristics. The sample consisted of recreationally resistance-trained individuals, and therefore extrapolation to highly trained strength athletes, clinical populations, or untrained individuals should be made with caution. Training history and baseline adaptive status are known to modulate both hypertrophic and molecular signaling responses to resistance exercise, which may constrain generalizability across populations. Finally, although a broad panel of anabolic and proteolytic signaling markers was assessed, dynamic measurements of muscle protein synthesis and breakdown rates were not performed. Such measures would provide a more direct index of net protein balance and could further clarify how different volume progression strategies influence muscle remodeling. This is particularly relevant given the known temporal dissociation between acute signaling protein activation and integrated myofibrillar protein synthesis responses (West, Kujbida et al. 2009).

## Conclusions

In conclusion, in resistance-trained individuals, an abrupt large increase in weekly resistance training volume (+120%) did not attenuate hypertrophy compared with a modest increase (+20%). Additionally, acute and chronic molecular markers related to anabolic regulation, proteolysis, and autophagy exhibited broadly similar responses across protocols. Collectively, these findings indicate that, within the present experimental context, resistance-trained skeletal muscle can accommodate a large abrupt increase in weekly set volume without detectable morphological impairment or systematic divergence in the molecular markers assessed.

## DATA AVAILABILITY

Source data are available from the corresponding authors upon reasonable request.

## FUNDING

This work was supported by The São Paulo Research Foundation (FAPESP, #2024/14207-0 to JBBC and #2024/12893-4 to JGAB), National Council for Scientific and Technological Development (#311387/2021-7 and #407748/2025-3 to C.A.L. and #140107/2023-1 to D.G.S.), and by laboratory funds from C.A.L. and M.D.R.

## CONFLICT OF INTEREST

The authors declare no conflicts of interest regarding the data presented in this study.

## Supplemental material

**Table 1.** Antibodies and associated IHC reagents used for experiments.

| Target | Host | Catalog Number | Dilution used | Manufacturer |
| --- | --- | --- | --- | --- |
| *Western blotting (base buffer was TBST + 5% bovine serum albumin)* |  |  |  |  |
| 20S proteasome core | Rabbit | BML-PW8155 | 1:1000 | Enzo |
| Calpain 1 | Rabbit | 2556 | 1:1000 | Cell Signaling Technology |
| Calpain 2 | Rabbit | 70655 | 1:1000 | Cell Signaling Technology |
| FOXO1 | Rabbit | 2880 | 1:1000 | Cell Signaling Technology |
| FOXO3 | Rabbit | 2497 | 1:1000 | Cell Signaling Technology |
| G3BP1 | Mouse | SC-365338 | 1:1000 | Santa Cruz Technology |
| LC3 | Rabbit | 2775 | 1:1000 | Cell Signaling Technology |
| p62 (SQSTM1) | Rabbit | 5114 | 1:1000 | Cell Signaling Technology |
| MyHCfrag | Mouse | A4.1025 | 1:200 | DSHB |
| Phospho-mTOR (Ser2448) | Rabbit | 5536 | 1:1000 | Cell Signaling Technology |
| mTOR | Rabbit | 2972 | 1:1000 | Cell Signaling Technology |
| Phospho-p70S6K (Thr389) | Rabbit | 2983 | 1:1000 | Cell Signaling Technology |
| p70S6K | Rabbit | 9202 | 1:1000 | Cell Signaling Technology |
| RPS6 | Rabbit | 2217 | 1:1000 | Cell Signaling Technology |
| Phospho-4EBP1 (Thr37/46) | Rabbit | 2855 | 1:1000 | Cell Signaling Technology |
| 4EBP1 | Rabbit | 9644 | 1:1000 | Cell Signaling Technology |
| SKIV2L2 | Mouse | SC-515828 | 1:1000 | Santa Cruz Technology |
| Ubiquitin | Rabbit | 3933 | 1:1000 | Cell Signaling Technology |
| Anti-rabbit HRP-conjugated | Goat | 7074 | 1:2000 | Cell Signaling Technology |
| Anti-mouse HRP-conjugated | Horse | 7076 | 1:2000 | Cell Signaling Technology |
| *Immunohistochemistry (base buffer was PBS)* |  |  |  |  |
| Dystrophin | Rabbit | GTX57970 | 1:100 | Genetex |
| MyHC I | Mouse (IgG2b) | BA-D5 | 1:100 | DSHB |
| Satellite cells | Mouse (IgG1) | PAX7s | 1:20 | DSHB |
| Nuclei (4,6-diamidino-2-phenylindole) | ----- | D3571 | 1:10,000 | Thermo Fisher |
| anti-rabbit IgG AF647 | Goat | A-21242 | 1:250 | Thermo Fisher |
| anti-mouse IgG2b Dylight488 | Goat | DI-1488 | 1:250 | Vector Laboratories |
| Anti-mouse IgG1 biotin (for SC amplification) | Goat |  | 1:1000 | Jackson ImmunoResearch |
| Streptavidin (for SC amplification) | ----- | S-911 | 1:500 | Thermo Fisher |
| Tyramide AF555 (for SC amplification) | ----- | B-40957 | 1:200 | Thermo Fisher |

Protein abbreviations: FOXO1, Forkhead box protein O1; FOXO3, Forkhead box protein O3; G3BP1, G3BP1 stress granule assembly factor 1; LC3, microtubule-associated protein 1B-light chain 3; p62, sequestosome 1; MyHCfrag: fragments of all myosin heavy chain isoforms; mTOR, mammalian target of rapamycin; p70S6K, ribosomal protein S6 kinase-1; RPS6, ribosomal protein S6; SKIV2L2, superkiller viralicidic activity 2-like 2. Other abbreviations: HRP, horseradish peroxidase; PBS, phosphate-buffered saline; phospho-, phosphorylated; SC, satellite cell; TBST, Tris-buffered saline with 0.1% Tween-20.

**Table 2.**
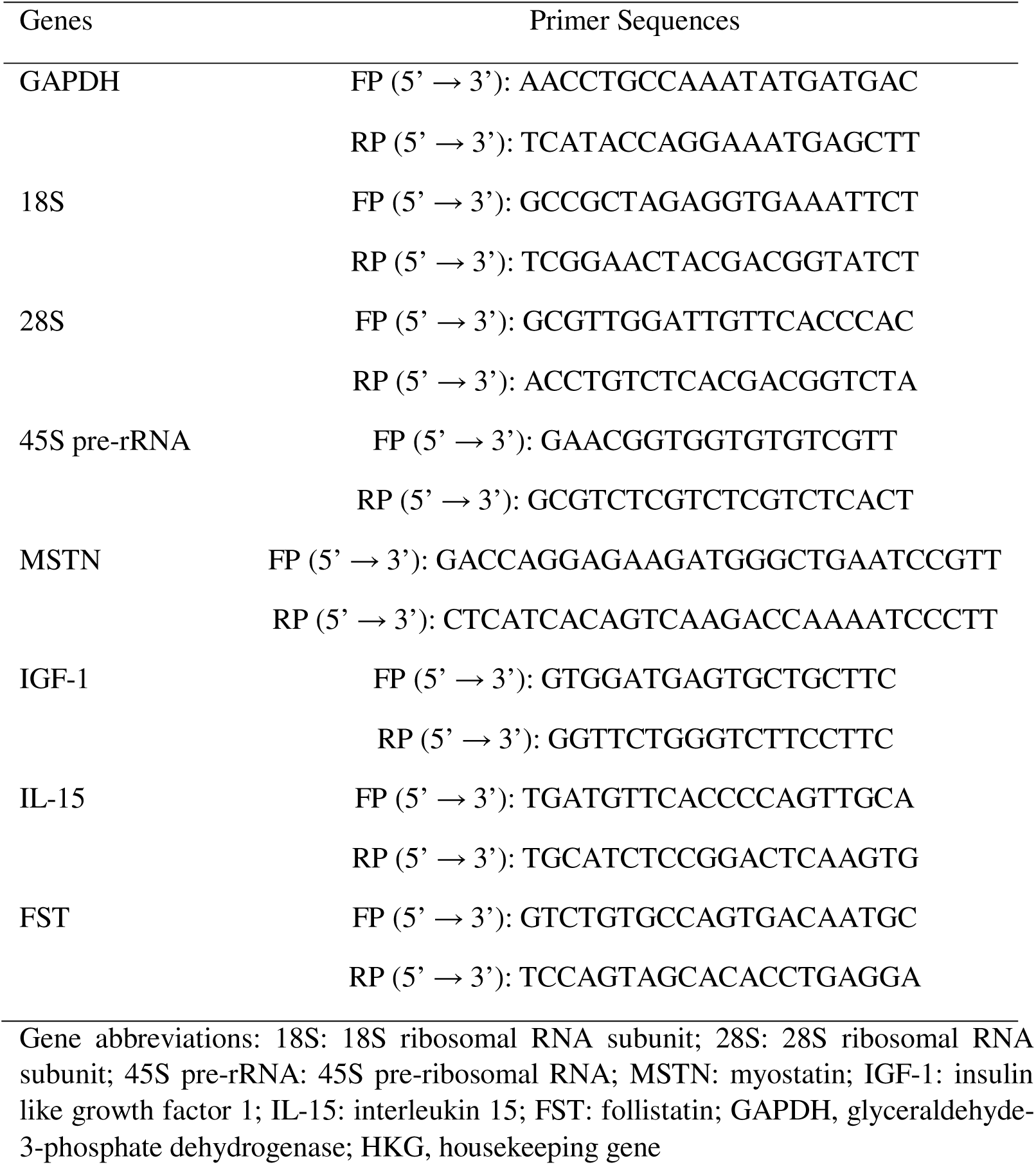
Primers used for qPCR experiments.

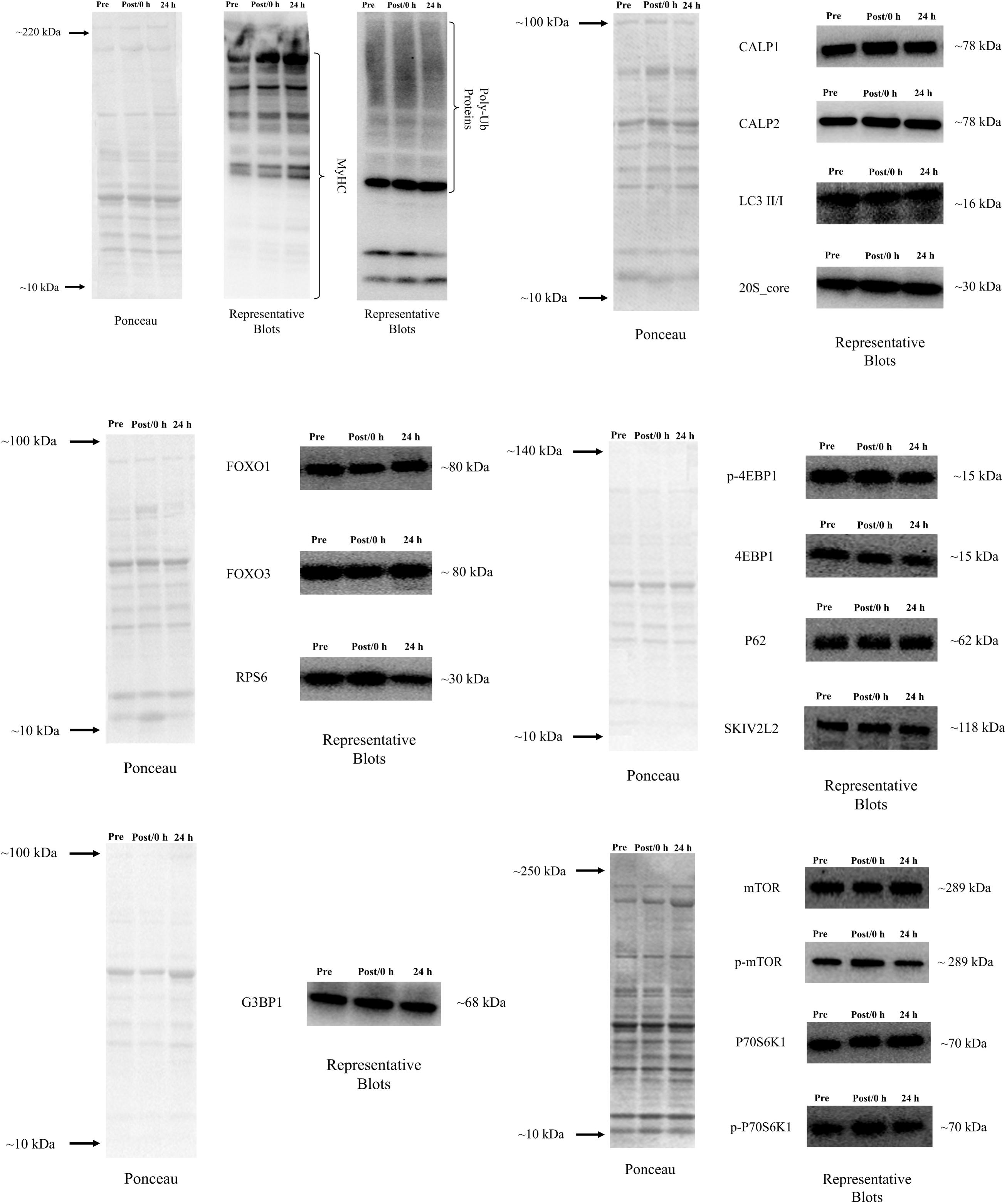

## Notes

### Competing Interest Statement

The authors have declared no competing interest.

